# A Competitive Fluorescence Immunoassay Based on Intermolecular Quenching Using an N-Terminally Fluorescent-Labeled IgG Antibody

**DOI:** 10.64898/2026.01.26.701636

**Authors:** Keisuke Fukunaga, Takayoshi Watanabe, Takahiro Hohsaka

**Affiliations:** School of Materials Science, Japan Advanced Institute of Science and Technology (JAIST), 1-1 Asahidai, Nomi, Ishikawa 923-1292, Japan; Institute for Tenure Track Promotion, University of Miyazaki, 1-1 Gakuen Kibanadai-Nishi, Miyazaki 889- 2192, Japan; MAQsys Inc., 3-2-1 Sakado, Takatsu, Kawasaki, Kanagawa 213-0012, Japan

**Author notes:** Corresponding author. (K.F.), (T.H.).

**Keywords:** Antibody, Immunoassay, FRET, Quencher, Thyroxine

## Abstract

The development of antibody-based fluorescent sensors relying on tryptophan-mediated quenching, such as Quenchbody (Q-body), often exhibits limited fluorescence responses because dye quenching depends on the location of tryptophan residues within the antibody. Here, we developed a competitive fluorescence immunoassay, termed an intermolecular Quenchbody (iQ-body), that utilizes intermolecular Förster resonance energy transfer (FRET) between an N-terminally fluorescent-labeled IgG antibody and an antigen-quencher conjugate. Anti-thyroxine IgG antibody (clone 6901 SPTN-5) showed minimal fluorescence changes upon antigen binding, despite N-terminal labeling with TAMRA. In contrast, the addition of an antigen-quencher conjugate (QSY9-T3) effectively quenched the TAMRA fluorescence via intermolecular FRET. Subsequent competitive displacement by thyroxine (T4) resulted in concentration-dependent fluorescence recovery. The iQ-body strategy provides a simple approach for constructing competitive fluorescence immunoassays for small-molecule targets using publicly available IgG antibodies.

## 1. Introduction

Rapid and straightforward detection of biomolecules is essential for applications in diagnostics, environmental monitoring, and biomedical research [1]. Although antibody-based assays are highly specific, conventional methods such as enzyme-linked immunosorbent assay (ELISA) [2] require multiple washing steps and long assay times, which limit their use in point-of-care and field applications. Consequently, “mix-and-read” sensing strategies [3], which enable direct signal transduction upon target binding, are highly desirable.

Quenchbody (Q-body) sensors [4-7] represent a class of fluorescent immunosensors based on antibody fragments whose fluorescence increases in response to antigen binding. Q-body typically consists of an antibody fragment, such as a single-chain variable fragment (scFv) [4,8], a Fab fragment [9], or a nanobody [10-12], that is site-specifically labeled with one or more fluorescent dyes in proximity to the antigen-binding site. In the absence of antigen, fluorescence is predominantly quenched via photoinduced electron transfer (PeT) from nearby intrinsic aromatic residues, particularly tryptophan [4,13,14]. Antigen binding triggers a conformational change of the antibody, which causes a dissociation of the fluorophore from hydrophobic patches on the antibody surface. This structural rearrangement relieves PeT-based quenching, resulting in a turn-on fluorescence signal that can be directly monitored in solution without washing steps. Q-bodies have demonstrated practical utility in a range of applications, including rapid detection of SARS-CoV-2 antigens [15], cytokines such as tumor necrosis factor-α (TNF-α) [16] and interleukin-6 (IL-6) [12], as well as environmental contaminants such as neonicotinoid pesticides [17].

In our previous study, we explored whether antibody-type fluorescent probes could be generated through site-specific fluorescent labeling of publicly available IgG monoclonal antibodies. We performed N-terminal-selective fluorescent labeling of IgG monoclonal antibodies, thereby converting them into antigen-responsive fluorescent antibody probes (**Fig. 1A**, successful example) [18]. This strategy successfully yielded antibody probes that exhibited fluorescence responses upon binding to their cognate antigens. However, this approach revealed several intrinsic limitations. First, the magnitude of the fluorescence intensity change (*F*/*F*0) strongly depended on the antibody clone. We hypothesize that the number and spatial distribution of tryptophan residue(s) within the antibody influence the fluorescence response, however detailed analysis was not feasible because the amino acid sequences of publicly available IgG antibodies are generally inaccessible. Second, many N-terminally fluorescent-labeled IgG antibodies failed to exhibit any fluorescence response upon antigen binding (data not shown) (**Fig. 1A**, unsuccessful example). In principle, effective fluorescence modulation requires tryptophan residue(s) positioned such that they can interact with the N-terminal fluorophore [12]; however, such favorable configurations are not universally present among IgG antibodies. As a result, extensive screening of monoclonal antibodies is required to identify suitable candidates for fluorescent probe construction. In addition, certain fluorescent dyes, such as cyanine, are inefficiently quenched by tryptophan residues [19]. This makes them incompatible with antibody-based fluorescent probes that rely on tryptophan-mediated quenching. Collectively, these limitations motivated us to develop an alternative strategy.

**Fig. 1.**
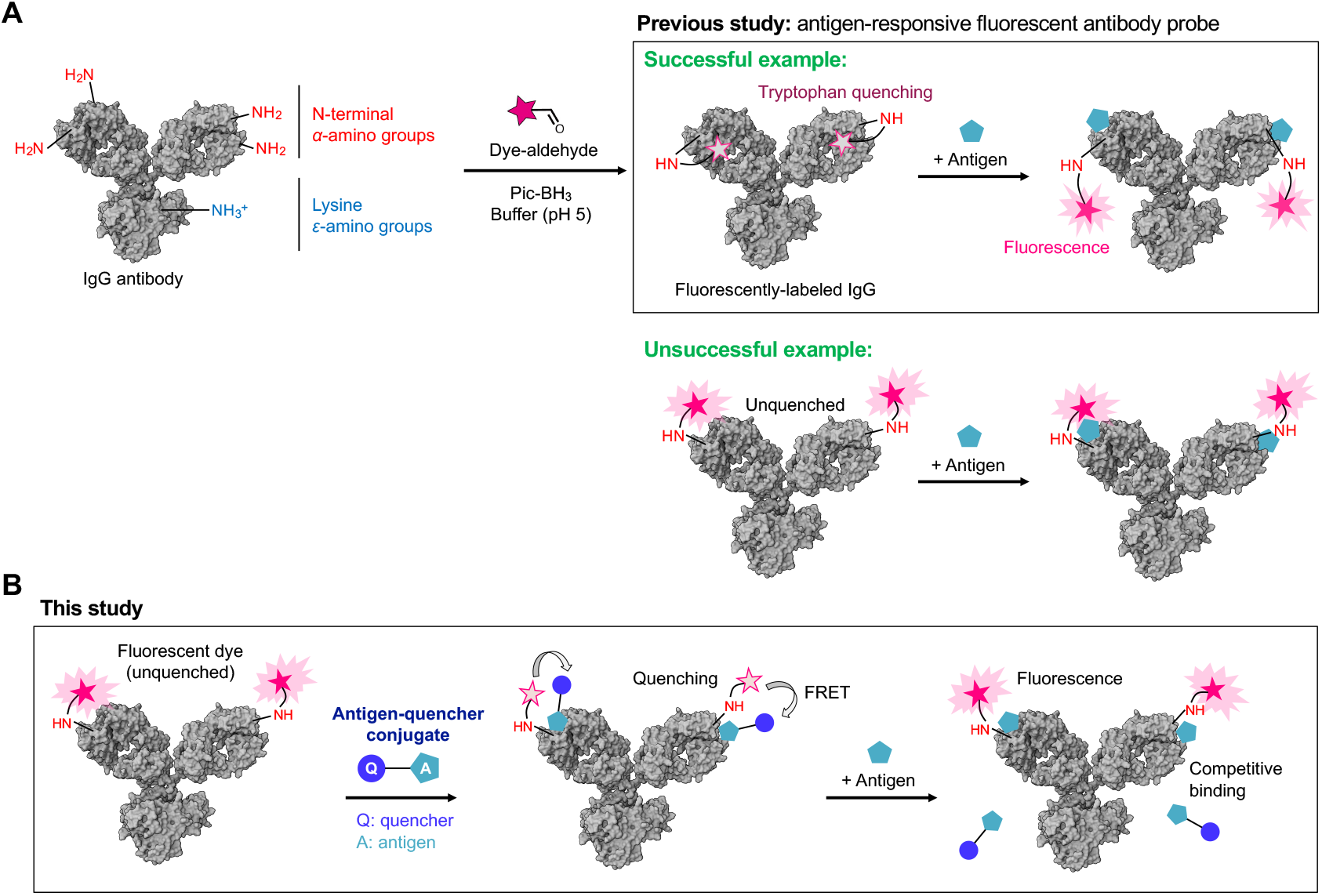
Schematic illustration of N-terminal-selective fluorescent labeling of IgG antibodies for the development of antigen-responsive probes. (A) A successful example represents a fluorescent antibody probe that exhibits a fluorescence response upon antigen binding. In contrast, an unsuccessful example represents a fluorescently-labeled antibody that does not exhibit a fluorescence response upon antigen binding. (B) Schematic illustration of the competitive fluorescence immunoassay using a pair of N-terminally fluorescent-labeled antibody and antigen-quencher conjugate. Pic-BH_3_: 2-picoline-borane. The IgG structure (PDB ID: 1IGY) [25] was generated using Mol* software [26].

To overcome the limitations associated with tryptophan-dependent quenching, we developed a competitive fluorescence immunoassay using a pair consisting of an N-terminally fluorescent-labeled IgG antibody and an antigen-quencher conjugate (intermolecular Quenchbody, hereafter referred to as iQ-body) (**Fig. 1B**). In this system, fluorescence quenching occurs via Förster resonance energy transfer (FRET) between the antibody-conjugated fluorophore and the antigen-quencher conjugate. The antigen-quencher conjugate quenches the fluorescence of the N-terminal fluorophore on the antibody, which is subsequently restored through competitive displacement by the free antigen, resulting in a dose-dependent turn-on signal. Fluorescence immunoassays based on intermolecular FRET were conceptually proposed decades ago [20] and have been applied to various haptens using antigen-quencher conjugates [21]. In previous intermolecular FRET-based immunoassays, random lysine labeling was employed, which can result in heterogeneous donor-acceptor distances. In contrast, N-terminal-selective fluorescent labeling provides more defined donor-acceptor distances in intermolecular FRET-based immunoassay. Unlike conventional lysine-based random labeling, which generates heterogeneous fluorophore modifications with variable spatial distributions, the N-terminal labeling restricts fluorophore attachment to a defined and structurally conserved position in IgG antibodies. Because the N-termini of the heavy and light chains are close to the antigen-binding site, the antigen-quencher conjugate is positioned at a well-defined distance from the antibody-bound fluorophore, resulting in a constrained FRET geometry. Another advantage of the iQ-body strategy is its potential compatibility with the conventional Q-body principle based on tryptophan-mediated quenching. By combining intermolecular FRET-based quenching with the intrinsic PeT between the fluorophore and tryptophan residues, an additive quenching effect could be achieved. Here, we report a competitive fluorescence immunoassay employing an N-terminally TAMRA-labeled anti-thyroxine IgG antibody with an antigen-quencher conjugate (QSY9-T3).

## 2. Materials and Methods

### 2.1. N-Terminal-Selective Fluorescent Labeling of Antibody

Fluorescent labeling of anti-thyroxine IgG antibody (clone 6901 SPTN-5, Medix Biochemica) was carried out in a 27 μL scale as previously described [18]. First, the antibody (approximately 9 μg) was buffer-exchanged into labeling buffer (50 mM sodium citrate, pH 4.8, 100 mM KCl, 0.1% polyethylene glycol 8000, and 0.05% Brij-35) by using an ultrafiltration device (30 kDa MWCO, PALL, Cat. No. OD030C34). The solution was adjusted to a volume of 21.4 μL and incubated on the ice for a few minutes in a 1.5 mL microtube. Subsequently, 1.1 μL of 1.92 mM 5(6)-TAMRA-X7-aldehyde (50% DMSO) was added. After mixing by tapping, 4.5 μL of 24 mM picoline borane (2.25% DMSO) was added. The 24 mM picoline borane solution was prepared just before use by diluting 1067 mM DMSO stock solution 45-fold with ultrapure water followed by vigorous vortexing. The labeling reaction was carried out at 4 ºC for 24 h. The resulting labeled antibody was purified twice by size-exclusion chromatography using gel filtration spin columns (EconoSpin, Gene Design, Cat. No. EP-31401), which were pre-equilibrated with equilibration buffer (20 mM sodium phosphate, pH 7.5, 100 mM NaCl, 0.1% polyethylene glycol 8000, and 0.05% Brij35).

### 2.2. SDS-PAGE and In-Gel Fluorescence Imaging

Protein samples were mixed with an equal volume of 2× sample buffer (Fujifilm-Wako, Cat. No. 193-11032) supplemented with 200 mM dithiothreitol and 10 mM ethylenediaminetetraacetic acid. The samples were denatured at 95 °C for 5 min immediately before electrophoresis. SDS-PAGE was performed using a laboratory-made 15% polyacrylamide gel. After the electrophoresis, in-gel fluorescence imaging was carried out using a fluoroimager (FMBIO-III, Hitachi Software Engineering). TAMRA was excited at 532 nm, and its emission was detected at 580 nm.

### 2.3. Fluorescence Spectroscopy

The TAMRA-labeled anti-thyroxine IgG antibody was 100-fold diluted with a measurement buffer (20 mM sodium phosphate, pH 7.5, 100 mM NaCl, 0.1% w/v polyethylene glycol 8000, and 0.005% Brij-35). After the addition of thyroxine (T4), the mixture was incubated at room temperature for several minutes. The mixture was transferred into a 5 × 5 mm quartz cuvette, and fluorescence spectra were recorded at 25 °C using a fluorescent spectrophotometer (Fluorolog-3, Horiba). TAMRA was excited at 550 nm, and emission spectra were acquired from 575 to 700 nm.

### 2.4. ELISA

Biotinylated thyroxine (biotin-X-T4) was synthesized in the previous study [18]. A protein G-coated 96-well plate (Thermo Fisher Scientific, Cat. No. 15131) was initially rinsed with PBS-B buffer (PBS supplemented with 0.05% Brij-35). IgG antibodies were diluted to 1 ng/μL in blocking buffer (PBS-B supplemented with 0.5% (w/v) bovine serum albumin), and 100 μL was added to each well. To immobilize the IgGs, the plate was incubated for 60 min at room temperature with shaking. After three washes with PBS-B, 100 μL of biotin-X-T4 was added, and then the plate was incubated for an additional 60 min with shaking. After three subsequent washes, alkaline phosphatase-conjugated streptavidin (Promega, Cat. No. V5591, 1:1000 dilution) was added and incubated for 30 min. After three washes, 100 μL of *p*-nitrophenyl phosphate (*p*NPP) solution (Thermo Fisher Scientific, Cat. No. 37621) was added. After 10 min incubation at room temperature with shaking, the absorbance at 405 nm was measured using a microplate reader (Multiskan GO, Thermo Fisher Scientific). Data analysis and curve fitting were performed using GraphPad Prism 10 software.

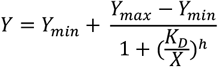

*Y*_*min*_: *minimum value of Y*

*Y*_*max*_: *maximum value of Y*

*K*_*D*_: *dissociation constant*

*X: molar concentration of antigen*

*h: Hill coefficient*

### 2.5. Synthesis of Antigen-Quencher Conjugate

QSY9-X-COOH: QSY9 succinimidyl ester (1 μmol, 80 μL of 12.5 mM DMSO solution) (Invitrogen, Cat. No. Q20131) and 6-aminohexanoic acid (5 μmol, 5 μL of 1 M aqueous solution) were mixed in a 1.5 mL microtube, followed by the addition of 0.1 M NaHCO_3_ (85 μL). After incubation at room temperature for 10 min with shaking, the reaction was quenched by the addition of 0.1% aqueous trifluoroacetic acid. The product was purified by preparative reverse-phase HPLC to afford QSY9-X-COOH (58% yield). After solvent evaporation, the residue was dissolved in DMSO. The concentration was determined by absorption at 575 nm using an extinction coefficient (*ε*) of 88,000 M^-1^ cm^-1^.

QSY9-X succinimidyl ester: QSY9-X-COOH (0.49 μmol, 63 μL of 7.12 mM DMSO solution) was mixed with 1-(3-dimethylaminopropyl)-3-ethylcarbodiimide hydrochloride (EDC-HCl) (4.41 μmol, 2.21 μL of 0.5 M DMSO solution), and *N*-hydroxysuccinimide (NHS) (4.41 μmol, 4.41 μL of 1 M DMSO solution). The mixture was incubated at 30 °C overnight with shaking. The resulting QSY9-X-SE was used for the subsequent coupling reaction without further purification.

QSY9-X-T3: QSY9-X-SE (0.1 μmol, 23.4 μL of 4.27 mM DMSO solution) and triiodothyronine (T3) (0.5 μmol of 10 mM DMSO solution) were mixed, followed by the addition of 100 mM aqueous NaHCO_3_ (73.4 μL). After 10 min incubation at room temperature, the reaction was quenched with 0.1% aqueous trifluoroacetic acid. The product was purified by preparative reverse-phase HPLC to afford QSY9-X-T3 (48% yield). The product was characterized by MALDI-TOF-MS (*m*/*z* calcd for [M+H]^+^ 1564.01, found 1564.01).

### 2.6. Competitive Fluorescence Immunoassay

For the quenching assay (**Fig. 3C**), the TAMRA-labeled anti-thyroxine IgG antibody was 50-fold diluted in the fluorescence measurement buffer. 1 μL of QSY9-X-T3 in DMSO was added to 50 μL of the antibody solution, and the mixture was incubated at room temperature for a few minutes. For the competitive assay (**Fig. 3D**), the antibody was 100-fold diluted in the same buffer, and QSY9-X-T3 was added at a final concentration of 48 nM, corresponding to the determined EC_50_ value of the antigen-quencher conjugate. And then, thyroxine (T4) (1 μL in DMSO) was added to 50 μL of this mixture and incubated at room temperature for several minutes.

A 50 μL aliquot of each sample was transferred into a microplate (Eppendorf, Cat. No. 30132700), which was subsequently sealed with a sealing film (Eppendorf, Cat. No. 0030132947). Fluorescence was measured using a real-time PCR system (Mx3005P, Agilent Technologies) with Cy3 channel (λ_ex_ = 545 nm, λ_em_ = 568 nm). Data were analyzed and plotted using GraphPad Prism 10 software.

## 3. Results and Discussions

### 3.1. Characterization of N-Terminally Fluorescent-Labeled Anti-Thyroxine IgG

The anti-thyroxine IgG antibody (clone 6901 SPTN-5) was selectively labeled at the N-termini with TAMRA via reductive amination (**Fig. 1A**). In-gel fluorescence imaging revealed the successful labeling of both the heavy and light chains (**Fig. 2A**). In antigen titration experiments, the TAMRA-labeled antibody exhibited a very small fluorescence response (maximum *F*/*F*_0_ = 1.15, the half-maximal effective concentration (EC_50_) = 2.3 nM) against its cognate antigen, thyroxine (T4) (**Fig. 2B**), indicating that this antibody clone does not inherently function as a conventional antigen-responsive fluorescent probe (i.e., Q-body). This finding is consistent with our previous results showing that antigen binding-dependent fluorescence recovery strongly depends on the presence and spatial arrangement of tryptophan residues near the fluorophore (data not shown). Next, we addressed whether the TAMRA-labeled IgG retains its ability to bind to antigen using enzyme-linked immunosorbent assay (ELISA) (**Fig. 2C**). The

**Fig. 2.**
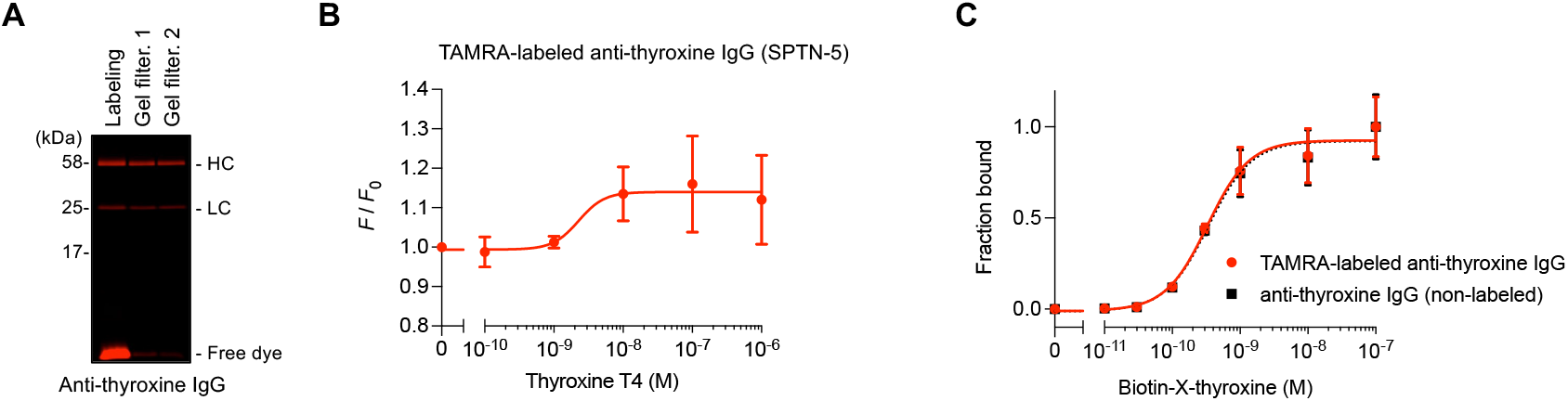
Synthesis and characterization of TAMRA-labeled anti-thyroxine IgG antibody (clone 6901 SPTN-5). (A) In-gel fluorescence imaging of TAMRA-labeled IgG. Proteins were resolved by 15% SDS-PAGE and visualized using a fluoroimager. HC: heavy chain; LC: light chain. (B) Antigen concentration-dependent fluorescence response of TAMRA-labeled anti-thyroxine IgG. Plots represent mean ± SD (*n* = 4). (C) Antigen-binding ability of non-labeled (black dotted line) and TAMRA-labeled (red solid line) antibodies by ELISA. Plots represent mean ± SD (*n* = 3).

ELISA analysis revealed that the TAMRA-labeled antibody binds to the antigen with an affinity that is comparable to that of the non-labeled antibody (dissociation constant (*K*_D_): 0.3 nM). The discrepancy between the observed EC_50_ and *K*_D_ values can be attributed to two factors. First, the fluorescence response requires an additional step, the displacement of the dye from the antibody surface, which may not occur immediately upon antigen binding, often resulting in an EC_50_ higher than *K*_D_ [22]. Second, under titration regime, where the concentration of the antibody probe exceeds the *K*_D_, the EC_50_ value depends on the probe concentration rather than reflecting the intrinsic binding affinity [23]. These results indicate that the antibody clone may be used in alternative fluorescence-based assay formats, while it is not suitable for direct signal-on sensing.

### 3.2. Design of a Competitive Fluorescence Immunoassay

The N-terminally TAMRA-labeled anti-thyroxine antibody was then applied to explore the possibility of intermolecular Q-body based on intermolecular quenching of TAMRA-labeled antibody by an external antigen-quencher conjugate (**Fig. 1B**). We synthesized a QSY9-labeled triiodothyronine (QSY9-X-T3) as the antigen-quencher conjugate (**Fig. 3A**). We selected QSY9 as a non-fluorescent quencher due to its strong spectral overlap with TAMRA emission (**Fig. 3B**), which enables efficient energy transfer upon antibody binding. Triiodothyronine (T3) was selected as the competitive antigen because its lower binding affinity than thyroxine (T4) facilitates efficient competitive displacement in the iQ-body assay. We evaluated the quenching effect of QSY9-X-T3 by antigen titration experiments (**Fig. 3C**). As expected, fluorescence intensity decreased in a dose-dependent manner, demonstrating effective quenching upon binding of the antigen-quencher conjugate. This result confirms that FRET occurs when the quencher is brought into proximity with the fluorophore via antigen–antibody interaction. The EC_50_ value of QSY9-X-T3 was determined to be approximately 48 nM. This concentration was used in subsequent competitive assays. Competitive binding assays were carried out by adding increasing concentrations of T4 in the presence of a fixed concentration of QSY9-X-T3 (48 nM) (**Fig. 3D**, red). Fluorescence intensity increased in a concentration-dependent manner (maximum *F*/*F*_0_ = 1.95, EC_50_ = 18 nM), indicating that T4 competitively displaced the antigen-quencher conjugate (i.e., QSY9-X-T3) from the antibody, thereby reducing FRET-based quenching. The observed EC_50_ value (18 nM) was higher than intrinsic *K*_D_ value of the antibody clone (0.3 nM) because of the competitive binding assay. In contrast, no fluorescence response was observed in the absence of QSY9-X-T3 (**Fig. 3D**, black). Taken together, these results demonstrate that antibodies unsuitable for direct signal-on sensing can be repurposed as effective probes in competitive fluorescence immunoassays.

**Fig. 3.**
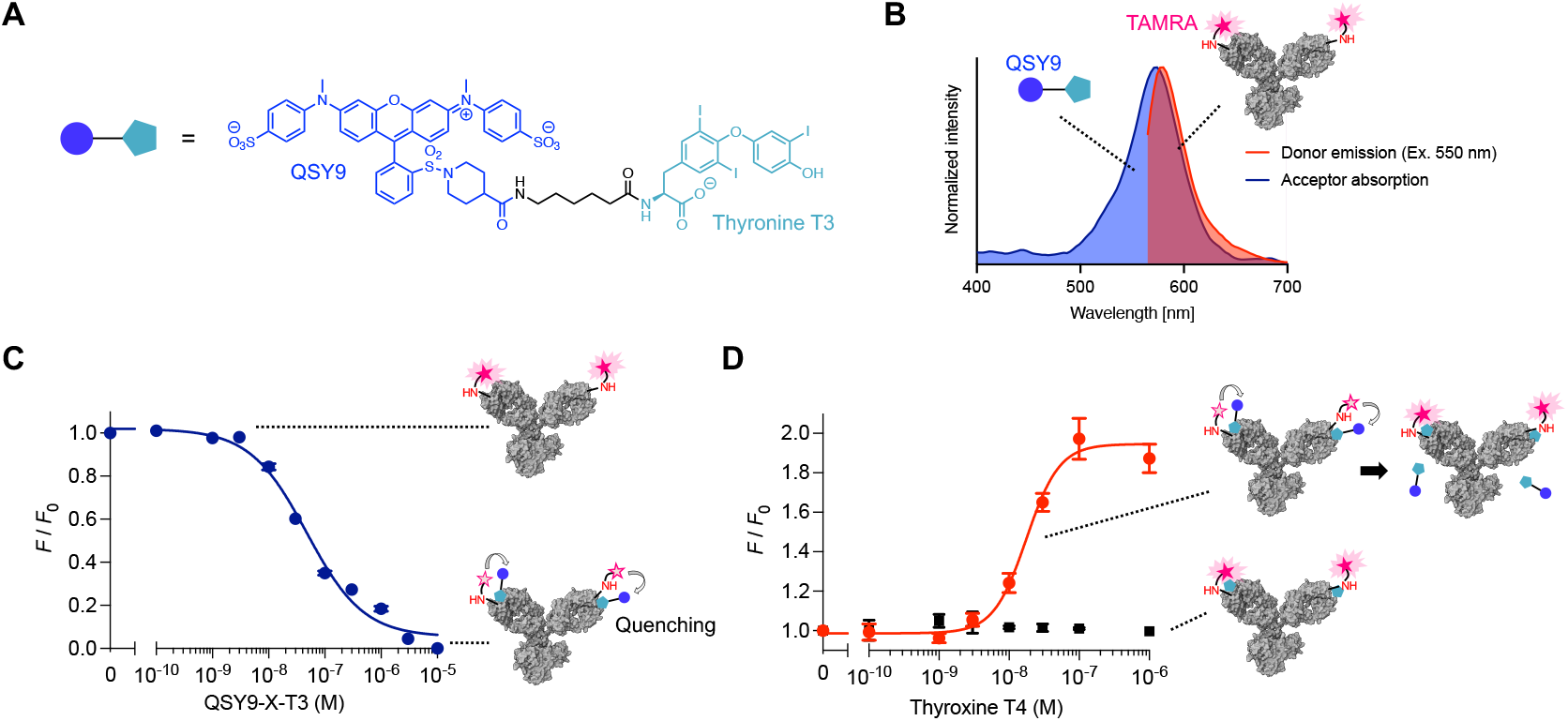
Competitive fluorescence immunoassay. (A) Chemical structure of QSY9-X-T3. (B) Spectral overlap between the acceptor (QSY9) absorption spectrum (blue) and the donor (TAMRA) emission spectrum (red). (C) Quenching of TAMRA-labeled anti-thyroxine IgG by QSY9-X-T3. (D) Competitive fluorescence response of the antibody to T4 antigen in the presence (red) or absence (black) of 48 nM QSY9-X-T3. Plots represent mean ± SD (*n* = 3).

### 3.3. Potential Applicability of Competitive Fluorescence Immunoassay

This competitive fluorescence immunoassay provides a complementary strategy to existing antibody-based fluorescent probes, including Q-body technology. Unlike conventional Q-body probes that rely on tryptophan-mediated quenching, the iQ-body does not require specific positioning of tryptophan residues within the antibody. In addition, the use of an external quencher can expand the repertoire of applicable fluorophores, including dyes that are inefficiently quenched by tryptophan. The assay operates in a homogeneous, mix-and-read format without washing steps, enabling rapid and straightforward detection. Because this system requires only an N-terminally fluorescent-labeled antibody and an appropriately designed antigen-quencher conjugate, it should be applicable to a wide range of small-molecule targets that are typically analyzed with competitive immunoassays [24]. Thus, the iQ-body strategy offers a versatile platform for constructing competitive fluorescence immunoassays.

## CRediT authorship contribution statement

**Keisuke Fukunaga:** Conceptualization; Funding Acquisition; Investigation; Formal Analysis; Visualization; Validation; Writing – Original Draft Preparation; Writing – Review & Editing. **Takayoshi Watanabe:** Resources; Investigation; Formal Analysis; Funding Acquisition; Writing – Review & Editing. **Takahiro Hohsaka:** Conceptualization; Funding Acquisition; Project Administration; Supervision; Writing – Review & Editing.

## Acknowledgements

We thank Dian Novitasari (JAIST) for her assistance with earlier studies.

